# Embryonic valproate exposure alters mesencephalic dopaminergic neurons distribution and septal dopaminergic gene expression in domestic chicks

**DOI:** 10.1101/2021.11.08.467690

**Authors:** Alice Adiletta, Alessandra Pross, Nicolò Taricco, Paola Sgadò

## Abstract

In recent years, the role of the dopaminergic system in the regulation of social behavior is being progressively outlined, and dysfunctions of the dopaminergic system are increasingly associated with neurodevelopmental disorders, including ASD. To further elucidate the role of the dopaminergic system in ASD, we investigated the effects of embryonic exposure to valproic acid (VPA) on the postnatal development of the mesencephalic DA system in the domestic chick. We found that VPA affected the rostro-caudal distribution of DA neurons, without changing the expression levels of several dopaminergic markers in the mesencephalon. We also investigated a potential consequence of this altered DA neuronal distribution in the septum, a social brain area previously associated to social behaviour in several vertebrate species, describing alterations in the expression of genes linked to DA neurotransmission. These findings support the emerging hypothesis of a role of DA dysfunction in ASD pathogenesis. Together with previous studies showing impairments of early social orienting behaviour, these data also support the use of the domestic chick model to investigate the neurobiological mechanisms involved in early ASD symptoms.

## Introduction

Autism spectrum disorder (ASD) comprise a group of neurodevelopmental conditions strongly characterized by impairments in sociability and social communication. In recent years, the role of the dopaminergic system in the regulation of social behaviour is being progressively outlined (Gunaydin and Deisseroth, 2014; Yamaguchi et al., 2017; Silva et al., 2020), often in association to motivational and reward mechanisms (Bariselli et al., 2018), and dysfunctions of the dopaminergic system in neurodevelopmental disorders and ASD has also been described (Scott-Van Zeeland et al., 2010; Supekar et al., 2018; Zürcher et al., 2021).

Approximately 75% of the total number of dopaminergic (DA) cells in the brain are located in the ventral part of the mesencephalon, giving rise to two major ascending DA systems that project to specific brain areas (Björklund and Dunnett, 2007). DA neurons in the ventral tegmental area (VTA) of the mesencephalon mainly project to the ventral striatum, the septum, the amygdala and the medial prefrontal cortex and constitute the mesocorticolimbic (or mesolimbic) pathway. Neurons of this pathway belong to the reward system and are thought to play a major role in controlling social reward, social learning and affiliative behaviour. On the other hand, neurons originating in the substantia nigra (SN) and innervating the dorsal striatum form the mesostriatal pathway, mainly involved in motor control. Despite the distinct categorization, neurons of the SN and the VTA have a complex organization and are intertwined in the mesencephalon, having also partially overlapping projections and sharing similar functions (Wise, 2009; Ilango et al., 2014). Furthermore, both DA mesencephalic nuclei are also blended with other type of neurons, as for example GABA and glutamate releasing neurons (Morales and Margolis, 2017) that contribute to their functional features (Bourdy et al., 2014).

In humans, the activation of the mesocorticolimbic pathway has been associated to processing of social stimuli (Spreckelmeyer et al., 2009), while a reduced activation of the same pathway has been described in children and adults with ASD (Scott-Van Zeeland et al., 2010; Supekar et al., 2018). Pharmacological studies in mice have shown that administration of DA mimetic and antagonizing drugs can modulate social interaction bidirectionally. In a seminal work Gunaydin and coworkers (Gunaydin et al., 2014) demonstrated a causal link between activation of VTA DA neurons and social interaction in mice using optogenetic tools. In addition, chemogenetic inhibition of VTA neurons projecting to Nucleus Accumbens (NAc) in mice has been associated to reduced social novelty seeking (Bariselli et al., 2018) and the molecular bases of such mechanisms in neurodevelopmental disorders are starting to be elucidated (Bariselli et al., 2016, 2018). Recent studies have also highlighted the role of the lateral septum, a limbic structure mainly innervated by the VTA DA neurons, in sociability and social novelty seeking in mice (Mesic et al., 2015), emphasizing the role of DA neurotransmission in the regulation of social behaviour (Hostetler et al., 2017; Shin et al., 2018). Overall, it appears that activation of mesolimbic system, connecting the VTA to NAc and to the LS, have a prosocial effect on pair and maternal bonding, social stimuli processing as well as on social interaction.

In this study, we harnessed the evolutionary conserved nature of the DA neuromodulatory system to investigate in domestic chicks the neurobiological mechanisms affected by VPA and potentially linked to the social behavioural deficits underlying ASD. VPA is an anticonvulsant known to interfere with development of the social brain, whose prenatal exposure is associated in humans with neural tube malformations, reduced cognitive function and an increased risk for developing ASD (Christensen et al., 2013). VPA embryonic exposure has been extensively used to model ASD core symptoms in diverse animal species (see for a review Bambini-Junior et al., 2014) including the domestic chick, where it induces alterations of several aspects of social behaviour (Nishigori et al., 2013; Sgadò et al., 2018; Lorenzi et al., 2019; Zachar et al., 2019; Adiletta et al., 2021). Here we examined the anatomical and molecular layout of the mesencephalic DA system in domestic chicks exposed to VPA during embryonic development. We found that VPA affected the rostro-caudal distribution of DA neurons, without changing the gene expression levels of several dopaminergic markers in the mesencephalon. We also investigated a potential consequence of this altered DA neuronal distribution in the septum, a social brain area previously associated to social behaviour in several vertebrate species (Lorenzi et al., 2017; Mayer et al., 2017; Clemens et al., 2020) and we observed alterations in the expression of genes linked to DA neurotransmission.

## Material and Methods

### Ethical approval

All experiments were conducted according to the current Italian and European Community laws for the ethical treatment of animals. The experimental procedures were approved by the Ethical Committee of the University of Trento and licensed by the Italian Health Ministry (permit number 986/2016-PR).

### Embryo injections

Fertilized eggs of domestic chicks (*Gallus gallus*), of the Ross 308 (Aviagen) strain, were obtained from a local commercial hatchery (Agricola Berica, Montegalda (VI), Italy). Upon arrival the eggs were incubated in the dark at 37.5 °C and 60% relative humidity, with rocking. One week before the predicted date of hatching, on embryonic day 14 (E14), fertilized eggs were selected by a light test and injected. Chick embryo injection was performed according to previous reports (Nishigori et al., 2013, Sgadò et al., 2018). Briefly, a small hole was made on the flat end of the egg and either VPA (Sodium Valproate, Sigma Aldrich, 35 μmoles) or vehicle (double distilled injectable water; CTRL group) were administered by dropping 200 μl of solution into the air sac of each fertilized egg. Eggs were assigned to the treatment groups randomly. After sealing the hole with paper tape, eggs were placed back in the incubator until E18, when they were prepared for hatching by incubation at 37.7 °C, with 72% humidity, and in complete darkness. The day of hatching was considered post-hatching day 0 (P0).

### Immunohistochemistry

After two days of dark incubation, P2 chicks were overdosed by an intramuscular injection of 0.05-ml per 10 g of body weight of Ketamine/Xylazine Solution (1:1 Ketamine 10 mg/ml + Xylazine 2 mg/ml). After 5 minutes chicks were transcardially perfused with phosphate-buffered saline (PBS) and ice-cold paraformaldehyde (4% PFA in PBS). Before removing the brain from the skulls, a coronal plane cut was performed using a stereotaxic apparatus (Kuenzel and Masson, 1988) to ensure correct orientation following the stereotaxic coordinates (45°) for coronal sectioning. Brains were then embedded in 7% bovine gelatine in PBS (4.2 g Bovine gelatine + 60 ml PBS at 40°C), using the plane cut to position the brain on the coronal plane. After cooling, the brains were post-fixed and cryopreserved in 4% PFA/PBS/20% sucrose for approximately 6 hours at room temperature, and then transferred to 30% Sucrose/0.4%PFA/PBS for further 72 h. Brains were then frozen in dry ice for 30 min before sectioning the entire brain in 60 μm coronal serial sections. For free-floating immunostaining, sections were washed 3 times in PBST (0,005% Triton/PBS) between each of the following steps. After incubation in 0.3% H_2_O_2_/PBS for 20 min, the sections were treated for 30 min with blocking solution (3% normal goat serum in PBS). Primary antibody reaction was carried out for 10 days at 4°C in blocking solution (1:1000 mouse monoclonal Tyrosine Hydroxylase antibody; T2928, Sigma-Aldrich). Sections were then incubated in biotinylated goat-anti-mouse antibody for 24 hours at 4 °C in blocking solution (1:200, BA-1000, Vector Laboratories). Color reaction was performed using the Vectastain Elite ABC Kit (PK-6100, Vector Laboratories) and the DAB kit (SK-4600, Vector Laboratories) following the manufacturer instructions.

### Cell counts

Counts of TH positive cells in the substantia nigra and ventral tegmental area were performed on 8 sets of serial sections per animal, sampled at 300µm intervals (n = 3 CTRL and 3 VPA chicks). Counting of TH immunoreactive cells was performed blind to the experimental condition using ZEN imaging software (Zeiss, Germany). All the sections were aligned on the rostral to caudal axis using Plate 34 of the chick brain atlas as reference (Puelles et al., 2007) and a rectangle of 150 × 250 pixels was placed over the samples and TH-positive cells within this area were manually counted. Cell densities were separately counted in the SN and the VTA as number of cells/area. For each analyzed brain slice, the sampling field was moved randomly through the area of interest at least four times for each dopaminergic subgroup.

### Tissue dissection

For microdissection of the SN and the VTA neurons, P2 chicks reared in complete darkness were euthanized via carbon dioxide gaseous asphyxiation, their brain extracted and then fast-frozen in dry-ice-cold isopentane solution. 100µm coronal sections were cut using a Leica CM1850 UV Cryostat at −15°C, and stored at −20°C. To better localize the targeted areas, sections were stained for 15 minutes with a 0.01% cresyl violet solution dissolved in 100% ethanol, and then progressively dehydrated in 75%, 90% and 100% ethanol (1 minute/each). All solutions were prepared fresh and filter-sterilized to avoid RNases contaminations. Substantia nigra and ventral tegmental area regions were finally dissected out using a 20G needle and immediately processed for total RNA extraction. For dissections of the septum, P2 chicks reared in darkness were euthanized by carbon dioxide gaseous asphyxiation, the brain extracted, and the area of interest directly dissected. Briefly, two coronal cuts were performed approximately 2 mm and 4 mm anterior to the bregma to isolate the septum in the anterior-posterior axis according to (Puelles et al., 2007) and the septum was carefully removed with forceps, fresh frozen in dry ice and immediately processed for total RNA extraction.

### Total RNA Extraction

Total RNA extraction from the septum was performed using the RNeasy Mini Kit (QIAGEN), while RNA from SN and VTA samples was extracted using the PicoPure™ RNA Isolation Kit (Applied Biosystems™, Thermo Fisher Scientific). Both extractions were performed according to the manufacturers’ instructions. Reverse transcription for both types of extracted materials was performed with the SuperScript™ VILO™ cDNA Synthesis Kit (Invitrogen, Thermo Fisher Scientific; Monza, Italy), following manufacturer’s instructions.

### Reverse Transcription-Quantitative Polymerase Chain Reaction (RT-qPCR)

RT-qPCR was carried out with PowerUp™ SYBR™ Green Master Mix (2x) (Thermo Fisher Scientific; Monza, Italy) for the septum and with SsoAdvanced™ Universal SYBR Green Supermix (Bio-Rad, Milan, Italy) for mesencephalic samples. Both reactions were performed using a CFX96™ Real-Time System (Bio-Rad, Milan, Italy). Commercially synthesized primers (Merck Life Science Srl, Milan, Italy) used in this work are listed in Table 1. Quantitation cycles (Cq) values were calculated using the second derivative maximum method. Data were normalized on the expression of TBP (TATA-Box Binding Protein) and HMBS (Hydroxymethylbilane Synthase) reference genes using the DeltaCt (dCt) method (Pfaffl, 2001).

**Table 1.**
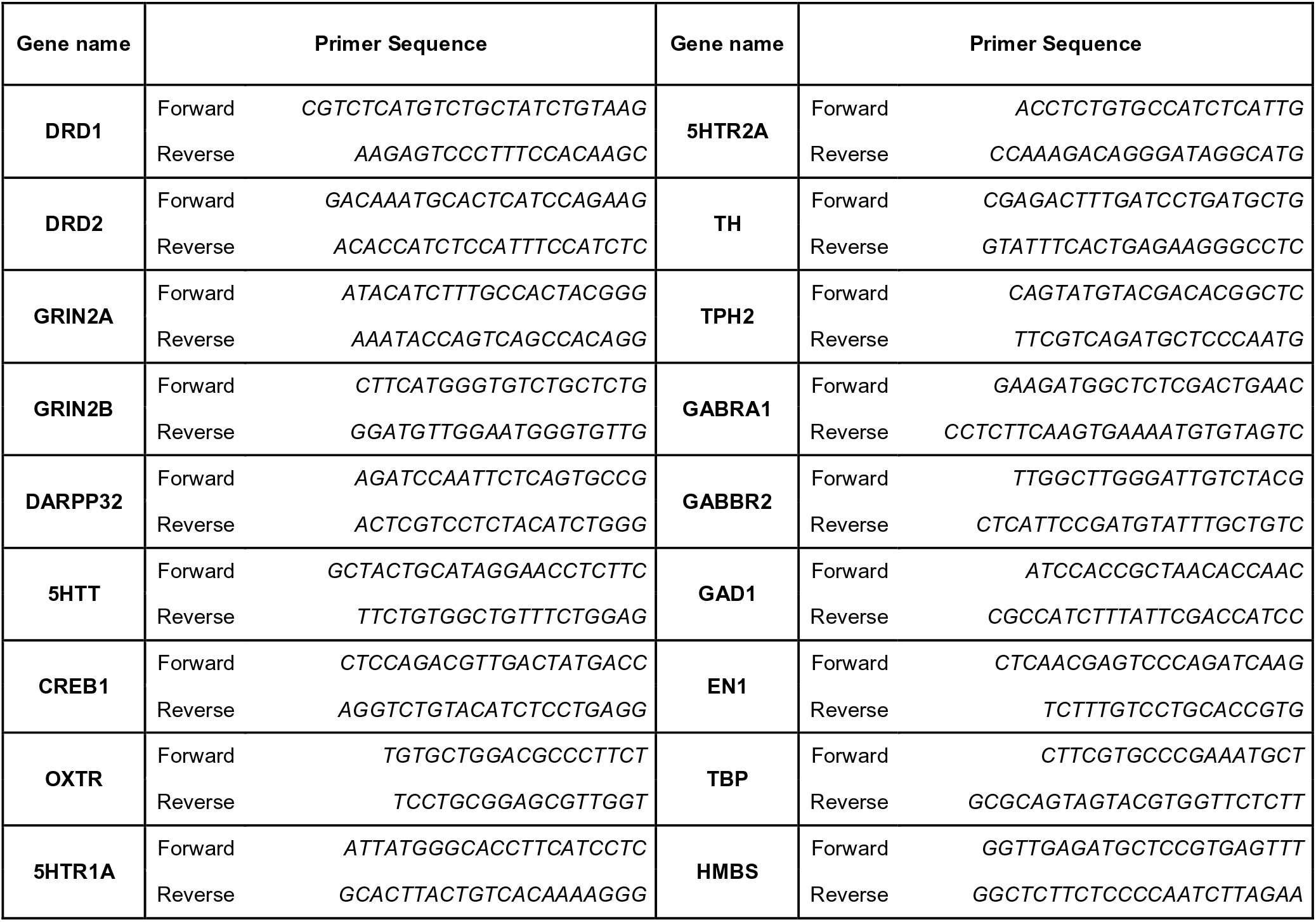
Primers used for RT-qPCR

### Statistical analysis

Statistical evaluation of the effect of treatment on the distribution and number of DA neurons of the different subgroups and of the log2 gene expression levels (dCt) was assessed using mixed-effect models using the nlme package in R (https://cran.r-project.org/web/packages/nlme/index.html). For Tukey pairwise comparison tests, we used the emmeans package in R (https://cran.r-project.org/web/packages/emmeans/index.html).

## Results

Previous studies demonstrated a remarkable effect of exposure to VPA during embryonic development on the dopaminergic system (Schiavi et al., 2019; Messina et al., 2020; Ádám et al., 2020; Román et al., 2021). To evaluate the effect of VPA on the dopaminergic system of domestic chicks, we performed immunohistochemical analysis and quantification of DA cell number in the SN and the VTA of VPA- and vehicle injected chicks, 48 hrs after hatching (P2). Brain sections from VPA- and vehicle-injected domestic chicks were immunolabeled for Tyrosine Hydroxylase (TH), the rate limiting enzyme for DA synthesis. Representative images of TH immunohistochemically labelled cells in the SN and VTA are shown in Fig1 A. TH-positive cells where then counted and DA cell densities (cells/area, see Methods) were quantified in each of the serial sections encompassing the mesencephalon, separating neurons of the SN from those of the VTA, and outlining their respective rostro-caudal distribution. To assess the effect of treatment (VPA and CTRL), DA group (SN and VTA) and rostro-caudal distribution (a set of 8 sections, from the most rostral to the most caudal) on DA density, we used a linear mixed model (LMM), considering treatment, group and section as fixed factors. We compared a model with random-intercepts-only to one with random slope and intercepts, without covariance between intercepts and slope, and found that the second model fitted the data significantly better. We found that the overall density of DA cells was significantly different between the two DA groups (SN vs VTA: F_(1,60)_ = 40.2371, p < 0.0001) and in the different rostro-caudal positions (sections: F_(7,60)_ = 10.5726, p < 0.0001) but no significant main effect of treatment was found in the overall number of DA cells (Fig. 1B; mean cells/area in substantia nigra CTRL 5.574 [95% C.I. 4.232 – 6.915], VPA 5.574 [95% C.I. 2.451 – 8.696] in ventral tegmental area CTRL 9.302 [95% C.I. 2.354 – 16.250], VPA 9.229 [95% C.I. 8.064 – 10.394; treatment: F_(1,4)_ = 0.7096, p = 0.4470). We also observed a significant interaction between treatment and section (treatment^*^section: F_(7,60)_ = 4.2912, p = 0.0007), indicating an effect of treatment on the rostro-caudal distribution of DA neurons. More interestingly, a triple interaction between treatment, section and group was observed (treatment^*^section^*^group: F_(7,60)_ = 2.2929, p = 0.0387), suggesting that the effect of treatment on the rostro-caudal distribution was different in the two DA subgroups (Fig. 1C and D). No other significant interactions were found between the other factors (treatment^*^group: F_(1, 60)_ = 0.0039, p = 0.9503; section^*^group: F_(7,60)_ = 0.50067, p = 0.4812). Given the differences on the overall density of DA cells in the subgroups, we analysed VTA and SN distribution separately. The pairwise comparison of the cell densities in VPA and vehicle-treated domestic chicks in each section of the separate groups (Fig. 1C and D) revealed a change in the distribution of the DA cell densities towards the posterior part of the mesencephalon, and thus a caudal shift in the distribution of DA neurons in VPA-injected chick compared to controls, more prominent in the substantia nigra (Fig 1C; CTRL vs VPA, section1 t_(4)_ = 8.7974, p = 0.0009; section 2 t_(4)_ = 3.1817, p = 0.0335; section 3 t_(4)_ = 2.4302, p = 0.0720; section 4 t_(4)_ = 1.9438, p = 0.1238; section 5 t_(4)_ = 0.5757, p = 0.5957; section 6 t_(4)_ = −3.5038, p = 0.0248, section 7 t_(4)_ = −3.7756, p = 0.0195; section 8 t_(4)_ = − 5.0059, p = 0.0075) than in the ventral tegmental area (Fig 1D; CTRL vs VPA, section1 t_(4)_ = 3.3448, p = 0.0287; section 2 t_(4)_ = 1.0207, p = 0.3651; section 3 t_(4)_ = 1.0597, p = 0.3490; section 4 t_(4)_ = −1.6494, p = 0.1744; section 5 t_(4)_ = −1.0495 p = 0.3532; section 6 t_(4)_ = 0.4148, p = 0.6996, section 7 t_(4)_ = −0.2410 p = 0.8214; section 8 t_(4)_ = −3.8945 p = 0.0176). Overall, our statistical analysis indicated a significant effect of VPA injection in the second embryonic week on the development of the mesencephalic dopaminergic neurons detectable at P2 as an alteration of the rostro-caudal distribution of DA cells.

**Figure 1.**
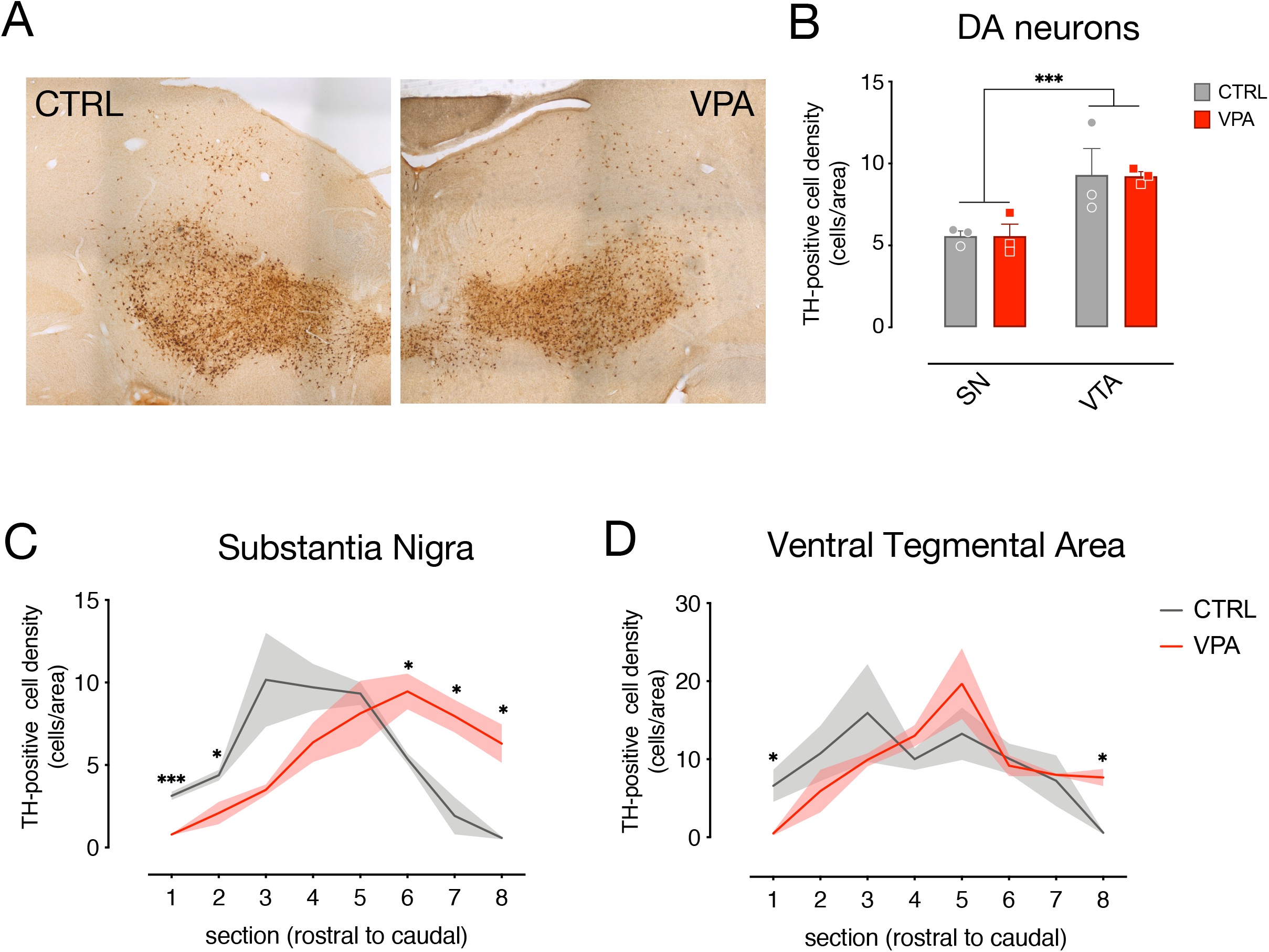
Immunohistochemical analysis and quantification of cell densities. (A) Brain sections from CTRL and VPA chicks immunolabeled for Tyrosine Hydroxylase (TH). (B) Number of TH-positive DA neurons in both SN and VTA. TH-positive cells’ count was performed on 8 sets of serial sections per animal, sampled at 300µm intervals. Rostro-caudal alignment of the brain sections was based on atlas reference (Plate 34 from Puelles et al., 2007). (C) Substantia Nigra DA cell density measured in its rostro-caudal distribution. (D) Ventral Tegmental Area DA cell density measured in its rostro-caudal distribution. ^*^ p < 0.05, ^***^ p < 0.001.

To assess the molecular changes induced by VPA on the dopaminergic system at P2, we micro-dissected DA neurons of the SN and VTA (Fig. 2A; n = 6 animal per treatment group, two independent experiments) in the entire rostro-caudal distribution, and measured the expression levels of genes involved in development (En1, TH, TPH2, Gad1) and neurotransmission (DRD1, DRD2, 5HTR1A, 5HTR2A, GABRA1, GABBR2). To assess the effect of treatment, DA group and transcripts, we used a linear mixed model, considering treatment, group and transcript as fixed factors and the experimental unit (experiment) as random factor. We compared a model with random-intercepts-only to one with random slopes and intercepts, and found that the random slopes and intercepts model fitted the data significantly better. The statistical analysis indicated a significant difference in the expression levels between the transcripts analysed in the two dopaminergic subgroups (transcripts: F_(8,152)_ = 406.6968, p < 0.0001; group: F_(1,152)_ = 3.7786, p = 0.0538; gene^*^group: F_(8,152)_ = 42.0532, p < 0.0001), however we could not detect any significant effect of VPA exposure at E14 on the expression of the genes at P2 (treatment: F_(1,10)_ = 0.0372, p = 0.8509; treatment^*^gene: F_(8,152)_ = 0.2526, p = 0.9795; treatment^*^group: F_(1,152)_ = 0.1096, p = 0.7410; treatment^*^gene^*^group: F_(8,152)_ = 0.7626, p = 0.6362). Independent on the treatment, some of the genes showed differences in expression levels between the two dopaminergic subgroups as indicated by the Tukey pairwise comparison for the transcripts in each DA subgroup (Fig. 2B; SN vs VTA). DRD2 (t_(152)_ = 4.8848, p < 0.0001), TH (t_(152)_ = 6.2522, p < 0.0001), GAD1 (t_(152)_ = 4.4819, p < 0.0001) and GABRA1 (t_(152)_ = 2.8885, p = 0.0044) show increased levels in the SN compared to the VTA, while TPH2 (t_(152)_ = − 15.5918, p < 0.0001) was expressed at higher levels in the VTA.

**Figure 2.**
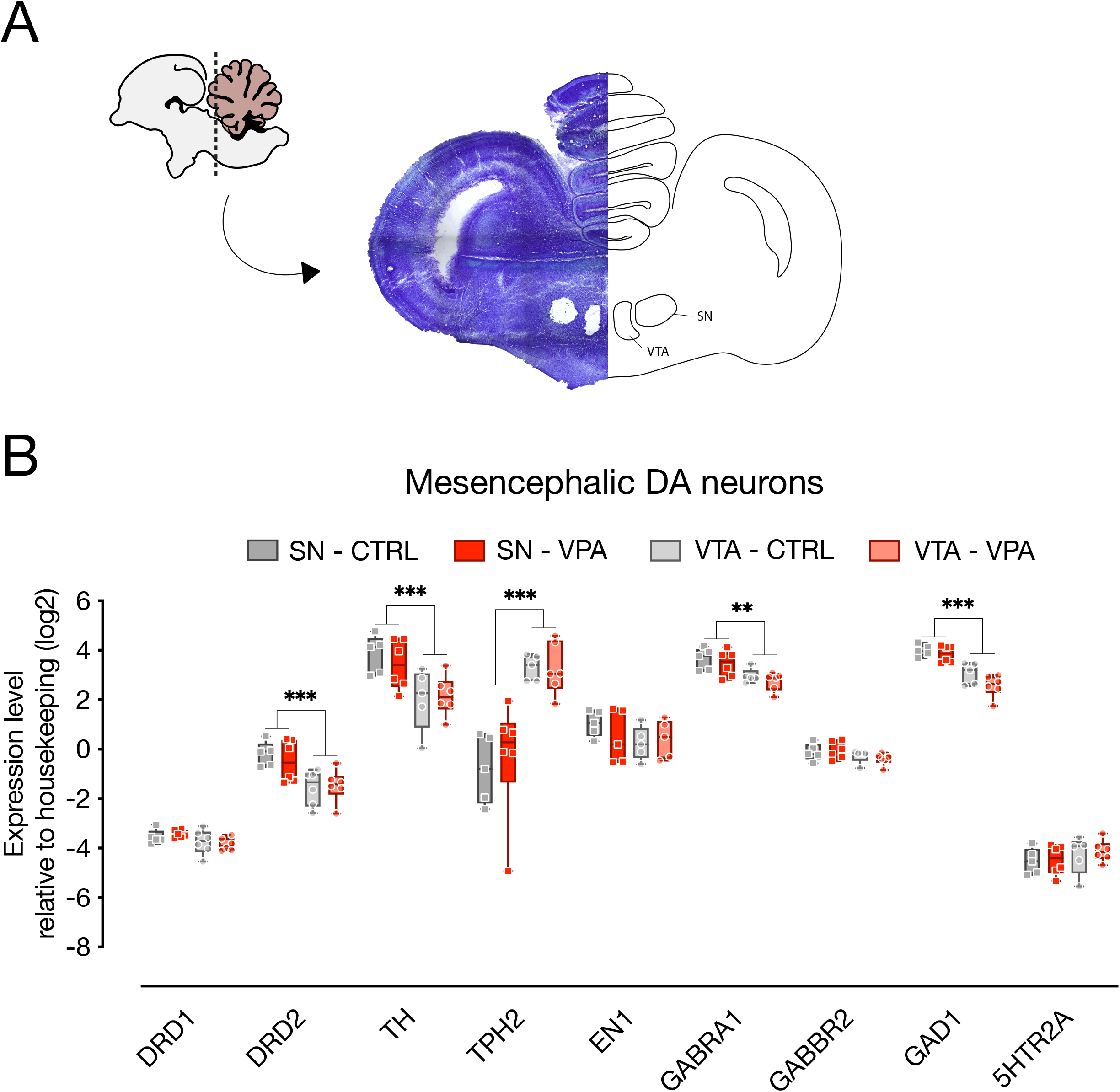
Gene expression levels in the SN and VTA. (A) Schematic representation of the microdissected areas of interest (SN, Substantia Nigra; VTA, Ventral Tegmental Area) from coronal sections (adapted from Plate 33 in Puelles et al., 2007). (B) Box and whisker plot (median, min to max) of relative expression (dCt, log2) values for each group. Changes in expression of DRD1, DRD2, TH, TPH2, EN1, GABRA1, GABBR2, GAD1 and 5HTR2A in both SN and VTA of VPA- and vehicle-injected chicks analyzed at P2. ^**^ p < 0.01, ^***^ p < 0.001.

We then examined gene expression changes in the septum, a brain area highly innervated by dopaminergic input coming from the SN and the VTA (Montagnese et al., 2008) and involved in social behaviour (Hostetler et al., 2017; Shin et al., 2018), that has been shown to activate in domestic chicks in response to exposure to conspecifics (Lorenzi et al., 2017; Mayer et al., 2017). We assessed the expression of gene involved in neurotransmission (Fig. 3; n = 8 animals per group, four independent experiments), like the dopamine receptors (DRD1 and DRD2) together with the DA responsive protein DARPP32, the serotonin receptors (5HTR1A and 5HTR2A) with the serotonin transporter 5HTT and genes involved in synaptic plasticity coding for the NMDA subunits (GRIN2A and GRIN2B), and CREB1. We also measured the expression levels of the mesotocin receptor (OXTR) the avian homolog of oxytocin, since the lateral part of the septum is known to receive consistent oxytocinergic innervation (Loveland et al., 2019; Horiai et al., 2020). We again used linear mixed models to evaluate the effect of treatment (CTRL vs VPA), DA subgroup (SN vs VPA) and transcripts, and found that the best fitting model was a random slopes and intercepts model with the same parameters used for analysis of gene expression in the mesencephalon. We found a significant main effect of treatment (F_(1,138)_ = 7.7599, p = 0.0061) and a significant differences in the levels of expression of the transcripts (F_(9,138)_ = 39.1627, p < 0.0001). We also observed a significant interaction between treatment and transcript (F_(9,138)_ = 5.1513, p < 0.0001), indicating an effect of treatment on some of the transcripts. The Tukey pairwise comparison indicated that expression of DRD1 (t_(138)_ = 2.0741, p = 0.0399), DARPP32 (t_(138)_ = 5.6749, p < 0.0001), and GRIN2A (t_(138)_ =2.9705, p = 0.0035) was decreased in VPA-injected chicks (Fig. 3), while 5HTT (t_(138)_ = −2.0297, p = 0.0443) expression was increased by the treatment, and expression of the other transcripts did not change (5HTR1A: t_(138)_ = 1.4917, p = 0.1381; 5HTR2A: t_(138)_ = −0.5621, p = 0.5749; CREB1: t_(138)_ = 0.7844, p = 0.4342; DRD2: t_(138)_ = 1.5030, p = 0.1351; GRIN2B: t_(138)_ = 1.6429, p = 0.1027; OXTR: t_(138)_ = −0.2558, p = 0.7985).

**Figure 3.**
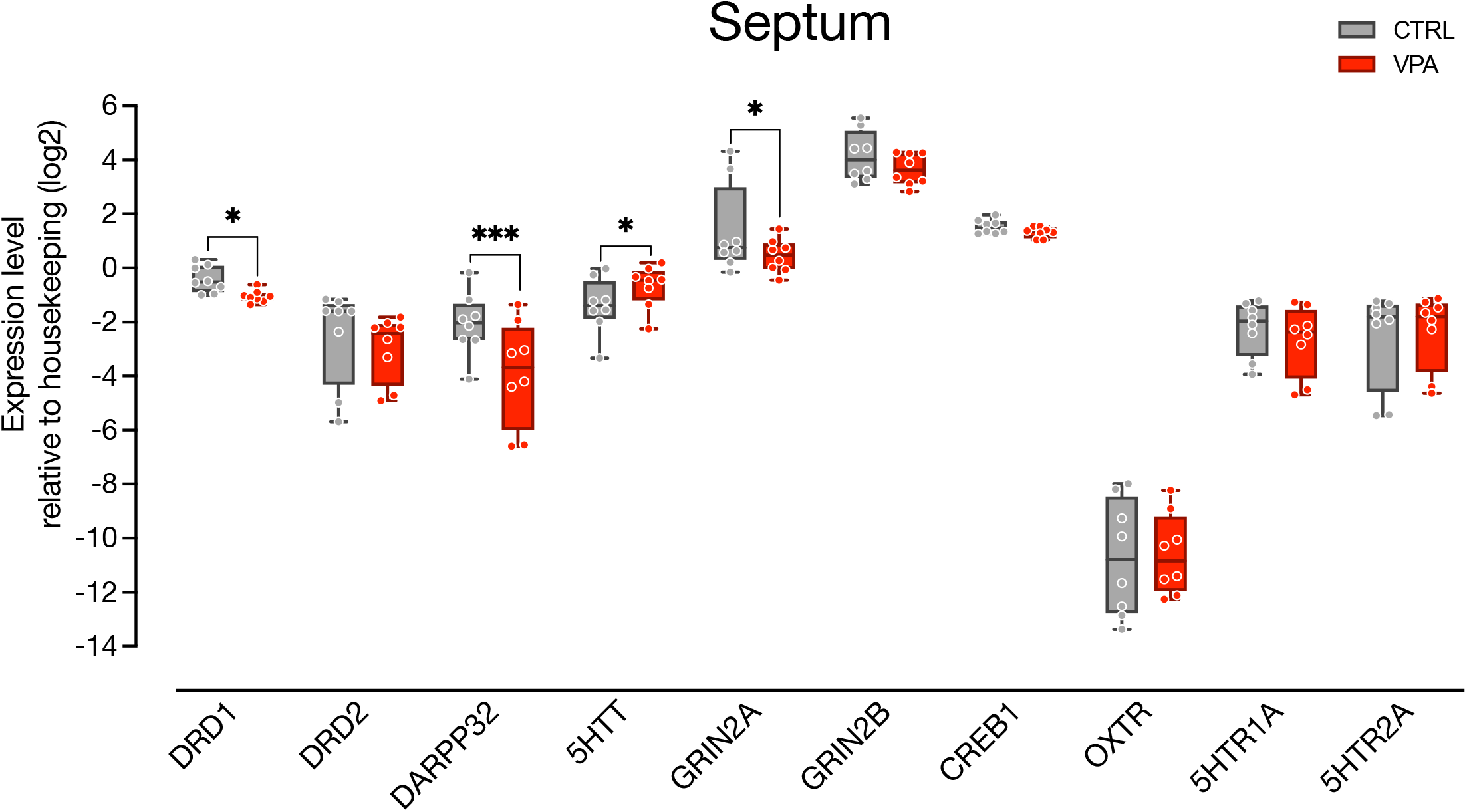
Gene expression levels in the septum. Box and whisker plot (median, min to max) of relative expression (dCt, log2) values for each group. Changes in expression of DRD1, DRD2, DARPP32, 5HTT, GRIN2A, GRIN2B, CREB1, OXTR, 5HTR1A and 5HTR2A in septum’s samples collected from P2 chicks embryonically exposed to VPA. Box and whiskers plot (median, min to max) of dCt values for each group. ^*^ p < 0.05, ^***^ p < 0.001.

## Discussion

Neuromodulatory systems, such as the those for the neurotransmitter dopamine, are evolutionarily very conserved, emerge in early embryonic development from ancient brain areas and are already mature at birth, innervating the neonatal brain. Thus, they represent an ideal target to modulate complex cognitive abilities originating in early development, such as social orienting behaviour. Notably, the dopaminergic system has been shown to influence key aspects of social and affiliative behaviours in humans and other vertebrates, and to cooperate in modulating key components of the social brain network (Gunaydin and Deisseroth, 2014). Moreover, accumulating evidence point to an involvement of DA neurotransmission in atypical social development in children with ASD (Scott-Van Zeeland et al., 2010; Supekar et al., 2018; Zürcher et al., 2021). In the present study, we investigated developmental dysregulations of the DA system in a VPA model of ASD implemented in domestic chicks, that could potentially contribute to the behavioural deficits observed in the chicks’ spontaneous responses to social stimuli, including altered response to faces (Adiletta et al., 2021). More specifically, we analyzed the number, distribution, and the developmental gene expression of DA neurons in the postnatal mesecephalon of domestic chicks embryonically exposed to VPA, and then assessed gene expression changes in the septum, a region of the social brain network highly innervated by DA terminals (Lindvall and Stenevi, 1978; Gaspar et al., 1985) and involved in sociability and social novelty seeking (Mesic et al., 2015).

Consistent with our hypothesis of an effect of VPA on the DA system, when sampling the rostro-caudal portions of the mesencephalon, we observed a caudal shift in the distribution of mesencephalic DA population at P2, nine days after embryonic exposure to VPA. Neuroanatomical alterations of mesencephalic TH population was already described in mice exposed to VPA by Ádám et al., (2020), however without analyses on the distribution of the DA population. Moreover, differently from the work of Ádám and colleagues (2020), here we found no difference in the total number of DA neurons (measured as number of cells/area) between the two treatments groups, suggesting the preservation of the overall profile of the DA population in our model, as indicated also by our gene expression analysis in the mesencephalon. Interestingly, VPA exposure was performed at E14, long after chick’s dopaminergic proliferation and differentiation events have terminated (Andersson et al., 2006), deeming unlikely a direct influence of VPA on neurogenesis or differentiation of new DA neurons. To investigate the potential consequences of the observed rostro-caudal shift in the distribution of the mesencephalic DA neurons, we have also assessed changes in the expression of genes involved in DA neurotransmission in one of the target regions of the mesocorticolimbic pathway, the septum. We found that DRD1, DARPP32 and GRIN2A were downregulated upon VPA exposure, consistent with deficits in dopaminergic signaling in this brain region. We also found that expression of the serotonin transporter (5HTT) was increased in the septum, suggestive of alterations also in serotonergic neurotransmission. Several studies and meta-analysis have confirmed an increase in 5HT in the blood (hyperserotonemia) of autistic individuals, such that hyperserotonemia has become a reliable biomarker for these disorders (see for a review Lam et al., 2006; and Gabriele et al., 2014). Epidemiological and animal model studies have suggested that perinatal alterations in 5HT, either above or below typical levels, may cause social behavioural deficits resembling ASD (Garbarino et al., 2019).

Previous studies have reported alterations in the number of DA neurons or in DA neurotransmission in several animal models of ASD (see for a recent review Kosillo and Bateup, 2021). A reduction in the number of DA neurons in the SN (but not the VTA) was found in adult mice lacking the Fmr1 gene (Fish et al., 2013). Further studies demonstrated reduced striatal DA transmission and striatal DA re-uptake without significant changes in the striatal tissue DA content in Fmr1 mutant mice (Fulks et al., 2010; Sørensen et al., 2015). Alterations in DA-mediated responses have also been reported in the BTBR mice (Squillace et al., 2014), a model for idiopathic autism, accompanied by decreased TH expression in several DA innervated brain regions (Chao et al., 2020). Interestingly, intranasal dopamine administration efficiently rescued the cognitive and social deficits of both BTBR and Fmr1 mutant ASD models (Chao et al., 2020), suggesting a causal role of DA deficiency in the behavioural phenotype of the mice.

Notably, in our study, we investigated for the first time DA-related deficits on an ASD model implemented in the domestic chicks. Differently from other common animal models, chicks are precocial species able to independently behave soon after hatching (Versace and Vallortigara, 2015), displaying remarkable early and spontaneous social responses, already shown to have similar features to the social orienting behaviour observed in human newborns (Di Giorgio et al., 2017). Domestic chicks also enable to study early neurodevelopmental mechanisms, without the interference of sophisticated, divergent, strategies of adaptive learning that emerge later. We thus believe that the domestic chick represents an ideal candidate model to study the causal relationship between social orienting behaviour, emerging at early postnatal stages, and any underlying neurobiological alterations mediated by VPA or any other genetic manipulation associated to ASD. Further studies should thus investigate the potential causal relationship between DA signaling alterations and the early social orienting deficits observed in VPA exposed chicks, including impairments in face processing and affiliative behaviour.

## Author Contributions

PS conceived and designed the experiments; AA and AP conducted the experiments; NT provided technical support; PS and AA analyzed the data; AA and PS wrote the manuscript. All authors read and approved the final manuscript.

## Acknowledgments

We thank Giorgio Vallortigara and Andrea Messina for their support. Dr. Tommaso Pecchia for help with the experimental apparatus, Grazia Gambardella and Roberta Guidolin for administrative help and Ciro Petrone for animal facility management.

## Data Availability Statement

All data generated or analysed during this study are included in this published article (see supplementary information).

